# Tanycyte-independent control of hypothalamic leptin signaling

**DOI:** 10.1101/528794

**Authors:** Sooyeon Yoo, David Cha, Dong Won Kim, Thanh V. Hoang, Seth Blackshaw

## Abstract

Leptin is secreted by adipocytes to regulate appetite and body weight. Recent studies have reported that tanycytes actively transport circulating leptin across the brain barrier into the hypothalamus, and are required for normal levels of hypothalamic leptin signaling. However, direct evidence for leptin receptor (*LepR*) expression is lacking, and the effect of tanycyte-specific deletion of *LepR* has not been investigated. In this study, we analyze the expression and function of the tanycytic *LepR* in mice. Using single-molecule fluorescent in situ hybridization (smfISH), RT-qPCR, single-cell RNA sequencing (scRNA-Seq), and selective deletion of the *LepR* in tanycytes, we are unable to detect expression of *LepR* in the tanycytes. Tanycyte-specific deletion of *LepR* likewise did not affect leptin-induced pSTAT3 expression in hypothalamic neurons, regardless of whether leptin was delivered by intraperitoneal or intracerebroventricular injection. Finally, we use activity-regulated scRNA-Seq (act-Seq) to comprehensively profile leptin-induced changes in gene expression in all cell types in mediobasal hypothalamus. Clear evidence for leptin signaling is only seen in endothelial cells and subsets of neurons, although virtually all cell types show leptin-induced changes in gene expression. We thus conclude that *LepR* expression in tanycytes is either absent or undetectably low, that tanycytes do not directly regulate hypothalamic leptin signaling through a *LepR*-dependent mechanism, and that leptin regulates gene expression in diverse hypothalamic cell types through both direct and indirect mechanisms.

## Introduction

Leptin is a cytokine-like hormone, mainly produced by adipocytes to signal levels of stored energy to the central nervous system (CNS). Hypothalamic neurons that regulate feeding and body weight are directly modulated by leptin, and show leptin-induced pSTAT3 expression within 30 minutes following either intraperitoneal (i.p.) or intracerebroventricular (i.c.v.) injection (Vaisse et al., 1996; Hubschle et al., 2001; Bates et al., 2003). However, the question of how circulating leptin is able to signal to hypothalamic neurons, which largely reside behind the blood-brain barrier (BBB), is still not fully resolved, despite many years of effort (El-Haschimi et al., 2000). Active transport of leptin into the hypothalamic parenchyma by endothelial cells has been put forward as one of the mechanisms (Banks et al., 1996; Di Spiezio et al., 2018). A second mechanism involving β2 tanycytes lining the median eminence of the hypothalamus has also been proposed (Langlet et al., 2013b; Balland et al., 2014); this model proposes that β2 tanycytes express leptin receptor (LepR), which binds circulating leptin. This in turn triggers transcytosis of the LepR-leptin complex, and transport of leptin into the lumen of the third ventricle, after which it is taken up into the hypothalamic parenchyma. Disruption of this process has been proposed as a possible mechanism behind the development of leptin resistance (Gao et al., 2014; Prevot et al., 2018). Increasing interest in the role of tanycytes has also raised the question of whether leptin signaling in tanycytes might directly affect regulation of hypothalamic metabolism more generally (Goodman and Hajihosseini, 2015; Djogo et al., 2016).

However, several key pieces of data on the potential physiological role of leptin signaling in tanycytes are missing. Biochemical approaches, which rely on antibody-based analysis of LepR expression and function (Hakansson et al., 1998; Mutze et al., 2006; Pan et al., 2008a), have been almost exclusively used to investigate this topic. These data, however, are dependent on the availability of highly specific antibodies, and recent work has clearly shown that many commercially available antibodies lack sufficient specificity, raising concerns about the accuracy of these results (Rhodes and Trimmer, 2006; Venkataraman et al., 2018). With this in mind, it is essential to determine whether *LepR* mRNA is actually expressed in tanycytes, and whether selective loss of function of *LepR* leads to disruption of leptin signaling in hypothalamus, before more substantial effort is invested in researching this topic.

In this study, we used a variety of highly sensitive techniques to investigate whether *LepR* mRNA is expressed in tanycytes, and to test whether leptin signaling in tanycytes is necessary for control of leptin signaling in hypothalamic neurons. Using a range of techniques – including single molecule fluorescent in situ hybridization (smfISH), quantitative PCR (RT-qPCR) of sorted tanycytes, and single-cell RNA sequencing (scRNA-Seq) analysis --we are unable to detect *LepR* mRNA expression in either adult or neonatal hypothalamic tanycytes, under conditions of either fasting or unrestricted food access. Moreover, selective deletion of *LepR* in tanycytes using the highly selective and efficient *Rax-CreER^T2^* line (Pak et al., 2014) fails to lead to any changes in pSTAT3 staining following either intraperitoneal or intracerebral delivery of recombinant leptin. Finally, act-Seq analysis of leptin-treated hypothalamus reveals that, while all hypothalamic cells showed some level of change in gene expression relative to saline-treated controls, substantial changes in known leptin-regulated genes are mainly observed in endothelial cells and subsets of neurons. These findings imply that tanycytes do not directly respond to leptin, and do not regulate leptin signaling in hypothalamic neurons via LepR.

## Materials and Methods

### Animals

*Rax-CreER*^*T*^*2*^^ mice generated in the laboratory (Pak et al., 2014) (JAX#025521) were bred with *LepR^lox/lox^* mice (Cohen et al., 2001) (JAX #008327) to generate tanycyte-specific LepR-KO mice. *Rax-CreER^T2^;Ai9 (R26-CAG-lsl-tdTom, JAX #007909)* and *Rax-CreER^T2^;CAG-Sun1/sfGFP* (Mo et al., 2015) (JAX #030952) were bred in the laboratory. To induce Cre recombination, tamoxifen was administered by either i.p. injection (1 mg, Sigma-Aldrich #H6278) at P28 for 3 consecutive days for fluorescent reporter expression or by feeding commercial tamoxifen-containing diet (EnvigoTeklad diets #TD.130856) for 3 weeks to delete *LepR* from tanycytes. *Rax-EGFP* BAC transgenic line (MMRRC #030564-UCD) was originally generated by the Gene Expression Nervous System Atlas Brain Atlas (GENSAT) Project (Gong et al., 2003). Seven weeks old C57BL/6 male mice were purchased from the Charles River Laboratories and used for scRNA-Seq analysis. All mice were housed in a climate-controlled pathogen free facility on a 14 hour-10 hour light/dark cycle (07:00 lights on – 19:00 lights off). All experimental procedures were pre-approved by the Institutional Animal Care and Use Committee (IACUC) of the Johns Hopkins University School of Medicine.

### Cell dissociation and FACS analysis

*Rax-CreER*^*T^2^*^;*CAG-Sun1/sfGFP* and *Rax-EGFP* BAC transgenic mice were used to isolate tanycytes using FACS. Briefly, tanycytes and nearby tissue regions were first micro-dissected from the adult brain using a chilled stainless steel brain matrix. Cells were dissociated using Papain Dissociation System (#LK003150, Worthington) following manufacturer’s instructions. Dissociated cells were resuspended in ice-cold PBS and flow-sorted into RLT lysis buffer (All Prep DNA/RNA micro Kit) using Sony SH800S Cell Sorter. Samples were stored at −80°C until RNA extraction.

### RNA extraction and RT-qPCR

RNA was extracted from both GFP-positive and GFP-negative cell fractions using AllPrep DNA/RNA micro Kit (#80284, Qiagen). For RT-qPCR, RNA samples were first reverse transcribed into cDNA using random primers and Superscript IV reverse transcriptase (#18091050, ThermoFisher) according to the manufacturer’s instructions. The qPCR assays were performed on the cDNA using GoTaq Green Master Mix (#M7122, Promega) using a StepOnePlus Real-time instrument (ThermoFisher). Intron-spanning primers were designed to specifically quantify targeted mRNA transcripts. Glyceraldehyde 3-phosphate dehydrogenase (*Gapdh*) expression was used as the endogenous control. *LepR* primers were designed to detect all transcript variants, including against the long form (*LepRb*), which is known to induce STAT3-mediated signaling, and the short forms (*LepRa* or *LepRc*), which were implicated in brain uptake of leptin. The following primers were used: *LepR*, Forward primer: GTGTCAGAAATTCTATGTGGTTTTG Reverse primer: TGGATATGCCAGGTTAAGTGC; *Crym*, Forward primer: GGCAACAGAGCCCATTTTAT Reverse primer: GTCATCCAGTTCTCGCCAGT; *Gapdh*, Forward primer: GACGTGCCGCCTGGAGAAAC Reverse primer: AGCCCAAGATGCCCTTCAGT. PCR specificity was monitored by determining the product melting temperature and by checking for the presence of a single DNA band on agarose gel analysis of the qRT-PCR products.

### Bulk RNA-sequencing of flow-sorted samples and bioinformatic analysis

Flow-sorted RNA samples from *Rax-EGFP* BAC transgenic mice were sent to the Deep Sequencing and Microarray Core (Johns Hopkins University) for library preparation and sequencing. Briefly, polyadenylated RNA was purified from the total RNA samples using Oligo dT conjugated magnetic beads and prepared for single-end sequencing according to the Illumina TruSeq RNA Sample Preparation Kit v2 (# RS-122-2001, Illumina). The libraries were sequenced for paired-end 75 cycles using the TruSeq SBS kit on NextSeq 500 system. Filtered sequencing reads were mapped to the mouse reference genome (mm10) using TopHat (Trapnell et al., 2009). FPKM value for each gene was estimated using Cufflink (Trapnell et al., 2012).

### Tissue processing, immunohistochemistry and RNAscope analysis

Mice were anesthetized with i.p injection of Tribromoethanol/Avertin and perfused transcardially with 1 x PBS followed by 2% PFA in 1x PBS. Brains were dissected and post-fixed in 2% PFA for overnight at 4°C. After washing, brains were incubated in 30% sucrose until brains sunk, then frozen in O.C.T. embedding compound. Brains were coronally sectioned at 25 µm thickness and stored in antifreeze solution at −20°C.

For immunohistochemistry, brain sections were post-fixed, if needed, and subjected to antigen retrieval treatment for pSTAT3 staining. Briefly, sections were sequentially incubated with 0.5% NaOH + 0.5% H_2_O_2_ for 20 min, 0.3% glycine for 10min, and 0.03% SDS for 10min, and blocked in 4% sheep serum/ 1% BSA/ 0.4% Triton X-100 in PBS for 1 hr at room temperature. Antibodies used were as follows: rabbit anti-pSTAT3 (1:1000, #9145, Cell Signaling Technology), mouse anti-HuC/D (1:200, #A-21271 Invitrogen), donkey anti-rabbit Alexa Fluor® 647 (1:500, #711-605-152, Jackson ImmunoResearch), goat anti-mouse IgG, Fcγ subclass 2b specific Alexa Fluor® 488 (1:500, #115-545-207, Jackson ImmunoResearch).

Sections were counterstained with DAPI and coverslipped using Vectashield antifade mounting medium (# H-1200, Vector Laboratories). All images were captured on a Zeiss LSM 700 Confocal at the Microscope Facility (Johns Hopkins University School of Medicine). All cell counts were performed blindly and manually on five or six sections per brain corresponding to −1.55mm, −1.67mm, −1.79mm, −1.91mm, −2.03mm, −2.15mm from Bregma. All pSTAT3 and HuC/D-double positive cell numbers counted and were normalized by the size (mm) of each hypothalamic nucleus measured using ImageJ. All values are expressed as means ± S.E.M. Comparison was analyzed by Student’s two-tailed t-test, and *P < 0.05* was considered statistically significant.

RNAscope was performed on 16 µm fixed (4% PFA in PBS) frozen adult sections. Sections were treated with 30% hydrogen peroxide for 10 minutes and then boiled (98-102^^^C) in 1X RNAscope target retrieval reagent for 15 minutes. Sections were next treated with RNAscope protease III at 40°C for 30 min. After protease incubation, sections were hybridized with *LepR* and *Rax* probes for 2 hours at 40°C. TSA Plus Cyanine 5 and fluorescein fluorophores were used to detect the hybridization signal. Sections were then counterstained with DAPI. Postnatal sections were not treated with retrieval reagent to preserve tissue morphology and were instead incubated in RNAscope Protease Plus at 40°C for 30 min.

### Leptin injection

Intracerebroventricular (i.c.v) administration of leptin was performed using cannulas (C315GS-5/SPC, 2.5 mm length of guide, Plastics One, Inc) implanted into the lateral ventricle (y: −0.3 mm, X: −1 mm, Z: 2.5 mm) of anaesthetized mice. Mice were given 1 week of recovery time following canulation and at 10 am, 2 µl of either aCSF (Tocris Bioscience #3525) or Leptin (0.5 µg/mice, Peprotech #450-31) was injected at 1 µl/min. Mice were returned to their to original cages for 1 hour, following which they were processed for act-Seq. For pSTAT3 analysis, mice were anaesthetized 30 min after i.c.v. injection or 5 min and 45 min after i.p. injection (3 mg/kg BW), and transcardially perfused with 2% PFA in PBS.

### Single-cell RNA-Seq library generation and analysis

Act-Seq (Wu et al., 2017) was performed based on previous method with a slight modification. Basically, brains were rapidly dissected and placed in cold 1x HBSS (Thermo Fisher Scientific, MA, USA) with Actinomycin D (3 µM, Sigma-Aldrich, MO, USA). 1 mm thick coronal slices (between Bregma –1.22 mm and −2.46 mm) were collected using adult mouse brain matrix (Kent Scientific, CT, USA). A juxtaventricular region that included portions of the dorsomedial hypothalamus, ventromedial hypothalamus, arcuate nucleus and the median eminence was then micro-dissected under a dissecting microscope. A total of 4 mice were used for each treatment group. Micro-dissected brain tissues were incubated in Hibernate-A media minus calcium (BrainBits LLC, IL, USA) with GlutaMAX (0.5 mM, Thermo Fisher Scientific), pronase (50 units/ml, Millipore Sigma), and Actinomycin D (15 µM), and were dissociated into single cells at 22’C with frequent agitation with fire-polished Pasteur pipette. Dissociated cells were filtered through 40 µM strainer and washed twice in Hibernate-A media with B-27 (2%, Thermo Fisher Scientific), GlutaMAX (0.5 mM), and Actinomycin D (3 µM). Cells were resuspended in this same media, with addition of RNase inhibitor (0.5 U/µl, Sigma-Aldrich).

Re-suspended cells were loaded into 10x Genomics Chromium Single Cell system (10x Genomics, CA, USA) using t v2 chemistry per manufacturer’s instructions, and libraries were sequenced on Illumina NextSeq with ~150 million reads per library. Sequencing results were processed through the Cell Ranger pipeline (10x Genomics) with default parameters. Both treatment groups (saline and leptin) were aggregated together for downstream analysis.

Seurat V2 (Butler et al., 2018) was used to perform downstream analysis following the standard pipeline, using cells with more than 500 genes and 1000 UMI counts. Clusters were annotated based on previous literature, and differential gene expression test was performed in each cluster, between treatment groups.

For analysis of scRNA-Seq data presented in Campbell *et al*, the gene expression matrix file was downloaded from the NCBI (accession number # GSE93374). Cells with > 800 expressed genes and genes expressed in >50 cells were used for the downstream analysis using Seurat V2. The tanycyte cell population was then subsetted computationally, and grouped into fasting and control samples.

## Results

### LepR expression is not detected in mature tanycytes

We used four distinct and complementary methods to measure *LepR* mRNA levels in tanycytes. We first performed RNA *in situ* hybridization in adult hypothalamic brain tissue using RNAscope, a highly sensitive and specific RNA visualization method. We successfully detected robust *Rax* mRNA expression in tanycytes (Figure 1A) and used it as a marker to distinguish any *LepR* expression in tanycytes from other cell types, including neurons, astrocytes and endothelial cells. As expected, substantial *LepR* mRNA expression was observed in hypothalamic parenchyma, in cells whose positions correspond to neurons of the arcuate nucleus. However, *LepR* mRNA expression is essentially undetectable in all *Rax*-positive tanycytes (Figure 1A).

**Figure 1.**
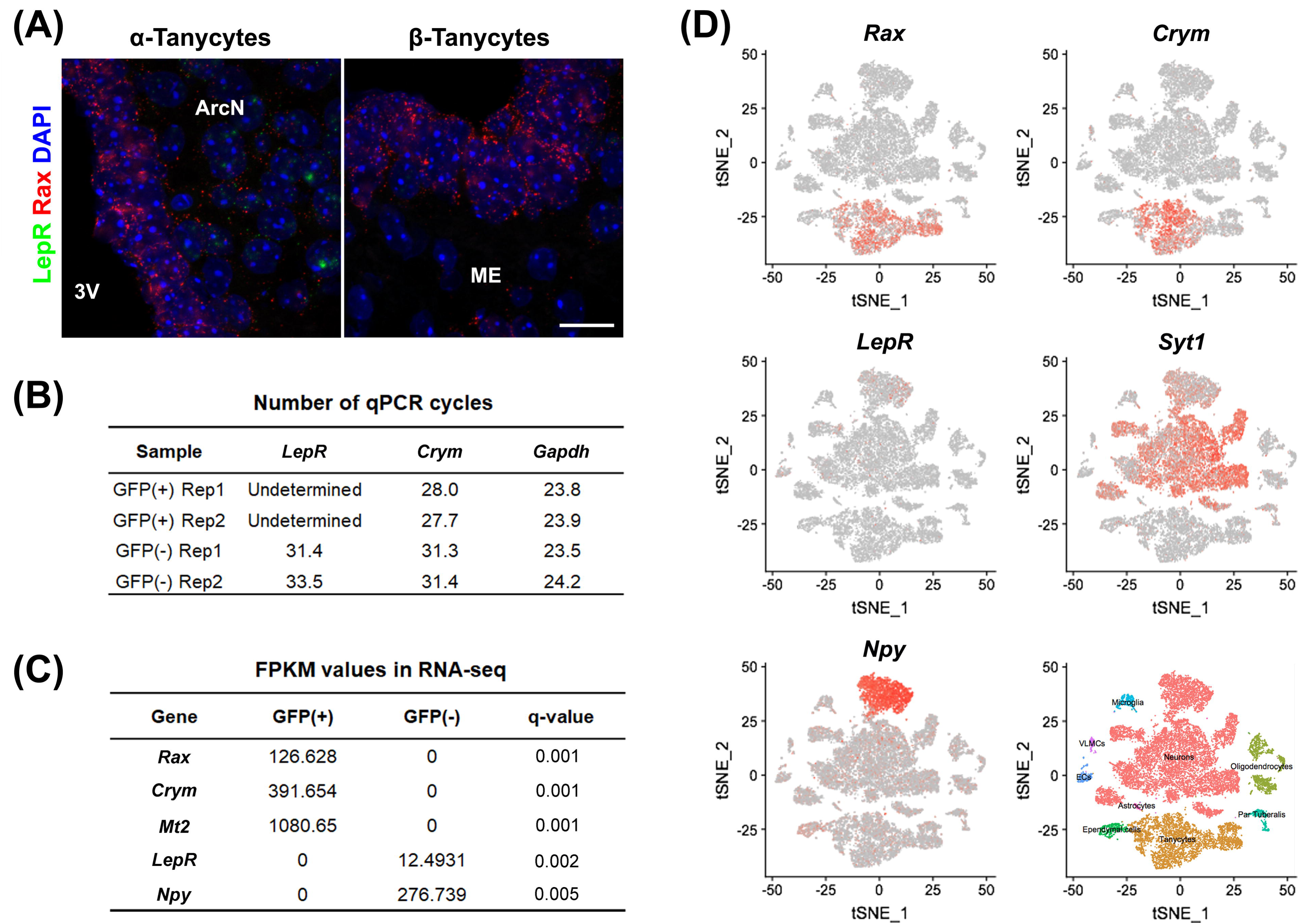
*LepR* mRNA expression is not detected in tanycytes of adult mice fed *ad libitum*. (A) Representative images for smfISH analysis using *Rax* (red) and *LepR* (green) probes showing α (left) and β tanycytes (right). (B) RT-qPCR analysis for LepR transcript in Rax-expressing tanycytes that were isolated from *Rax-CreER^T2^/lsl-Sun1-GFP* mice. *Crym* is a tanycyte marker. *Gapdh* expression, used as a loading control, showed similar abundance in each sample. (C) Differential expression analysis between GFP-positive and GFP-negative population sorted from *Rax-EGFP* mice displaying examples of highly enriched in each fraction and LepR. (D) tSNE plots using previously reported data from 20,921 cells isolated from the ArcN-ME. *Rax* and *Crym* were strongly expressed in the tanycyte cluster, and *LepR* in the *Syt1*-positive neuronal cluster, particularly in *Npy*-positive neurons. Scale bar: 20um (A).

We next tried to detect the *LepR* transcript in GFP-positive cells sorted by fluorescence-activated cell sorting (FACS) from *Rax-CreER^T2^;CAG-lsl-Sun1-GFP* mice (Mo et al., 2015). Tamoxifen injection induced Cre recombinase activation in *Rax*-expressing tanycytes, resulting in GFP expression restricted to the nuclear membrane. To confirm that GFP-positive sorted cells were indeed tanycytes, RT-qPCR analysis was performed on the tanycyte marker gene *Crym* (Campbell et al., 2017), and highly enriched *Crym* expression is observed in GFP-positive cells. *LepR* mRNA expression, however, is not detected the GFP-positive fraction, but is readily detected in the GFP-negative, tanycyte-depleted fraction (Figure 1B).

Next, we investigated whether *LepR* mRNA could be detected in tanycytes in bulk RNA-Seq data. Hypothalamic tissues from *Rax-EGFP* mice (Gong et al., 2003) were used for FACS sorting, and separated into GFP-positive and GFP-negative populations. Differential gene expression analysis of RNA-Seq data show a strong enrichment for tanycytes in the GFP-positive fraction, as confirmed by expression of tanycyte markers such as *Rax, Crym* and *Mt2*, and negligible contamination is detected from parenchymal *Npy*-expressing neurons. Consistent with RNAscope and RT-qPCR data, *LepR* transcript is not detected in GFP-positive tanycytes, but only found in GFP-negative cells (Figure 1C).

We also analyzed a recently published single cell RNA-Seq dataset from 20,921 ArcN-ME dissociated cells (Campbell et al., 2017). Cells are clustered into distinct cell types. The data show that the tanycyte cluster was clearly separated from other cell types, as confirmed by *Rax* and *Crym* expression (Figure 1D). In agreement with our data, scRNA-Seq showed that *LepR* expression was detected other cell types, notably in *Agrp/Npy*-positive neurons, but not in the tanycyte cluster.

### Fasting does not induce LepR mRNA in tanycytes

The previous experiments were performed in mice given *ad libitum* access to chow, and thus does not exclude that alternative dietary conditions might induce *LepR* expression in tanycytes. Fasting reduces circulating leptin levels, and increases *LepR* transcription in both *Pomc*-and *Agrp*-positive neurons (Baskin et al., 1998; Bennett et al., 1998; Mitchell et al., 2009). To determine whether this was also the case in tanycytes, we conducted smfISH on animals that had been fasted for 24 hours. As previously reported, *LepR* expression is substantially increased cells in hypothalamic parenchyma under these conditions. However, fasting does not induce *LepR* expression in tanycytes (Figure 2A). Consistent with this result, scRNA-Seq data from both fasted and control mice shows that *LepR* expression was barely detected in the tanycyte cluster under either condition, as marked by tanycyte-specific genes: *Col23a1* and *Slc16a2* (Figure 2C), while the cell number and level of expression is substantially increased in the neuronal cluster (*Syt1*-positive) following fasting, with greatest increases observed in *Npy/Agrp*-positive neurons (Figure 2D). Consistent with the previous report, an increased *Npy* and *Agrp* expression was observed in this subset of neurons after fasting (Figure 2D)(Korner et al., 2001). Taken together, the fact that we did not detect *LepR* expression in tanycytes using any of these techniques raises serious questions about potential physiological relevance of leptin signaling in tanycytes.

**Figure 2.**
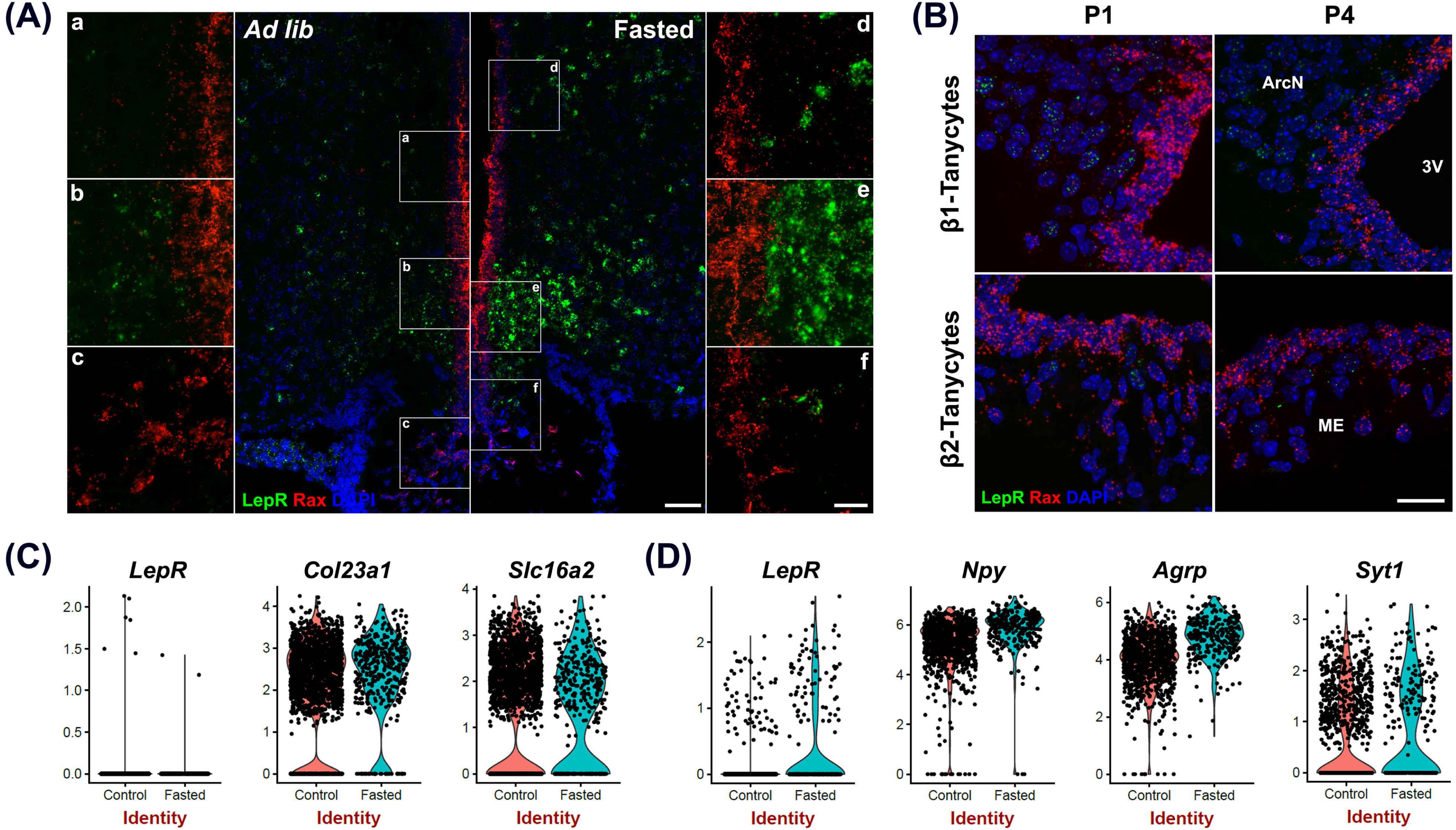
*LepR* mRNA expression is not detected in tanycytes of neonatal or fasted adult mice. (A) smfISH analysis using *Rax*(red) and *LepR*(green) probes in *ad lib* and fasted animals. (a-f) are the higher magnification images of the boxed area in (A). *Rax*-expressing red fluorescence labeled region represents the tanycytic layer. (B) Violin plots of *LepR* and known tanycytic markers *Col23a1* and *Slc16a2* in tanycyte cluster comparing between ad lib and fasted conditions. (C) Violin plots of *LepR* and known neuronal marker genes in *Npy*-positive cluster. (D) smfISH analysis using *Rax*(red) and *LepR*(green) probes in P1 and P4 mice. The upper panel shows β1 tanycytes and the lower panel shows β2 tanycytes. Scale bar: 50um (A), 20um (B), 20um (a-f)

### LepR mRNA was not detected in immature tanycytes

Previous studies using radioactive *in situ* hybridization reported *LepR* expression in neonatal rats in the ventricular zone of the arcuate nucleus, with expression emerging only later in parenchymal neurons (Cottrell et al., 2009), although *LepR* expression at this early stage does not appear to be associated with the Leptin transport across the blood-brain barrier (Pan et al., 2008b). To determine if the *LepR* is expressed in immature tanycytes in mice, we performed smfISH analysis at postnatal day (P)1 and P4. At both P1 and P4, *LepR* expression is accumulated in the parenchymal cells, presumably neurons, but is absent in tanycytic layer, as marked by *Rax* expression (Figure 2B).

### Leptin signaling in hypothalamic parenchyma is maintained following tanycyte-specific deletion of LepR

These findings still do not formally exclude a physiological role for very low levels of *LepR* expression in tanycytes. To directly address this, we crossed *Rax-CreER^T2^;R26-lsl-tdTom* with *LepR^lox/lox^* mice (Cohen et al., 2001) to generate *Rax-CreER^T2^;LepR^lox/lox^;CAG-lsl-tdTom* mice. *Cre*-dependent excision of the first coding exon of *LepR* in this mouse will lead to a null mutation that will disrupt function of all known *LepR* splicing variants. We previously observed tanycyte-specific Cre recombination using *Rax-CreER^T2^;CAG-lsl-tdTom* mice (Pak et al., 2014), and observe robust tdTomato expression in tanycytes following three weeks of administration of tamoxifen-infused chow (data not shown). SmfISH analysis does not detect *LepR* mRNA expression in the tanycytes from either the control or the *LepR* knockout mice by (Figure 3).

**Figure 3.**
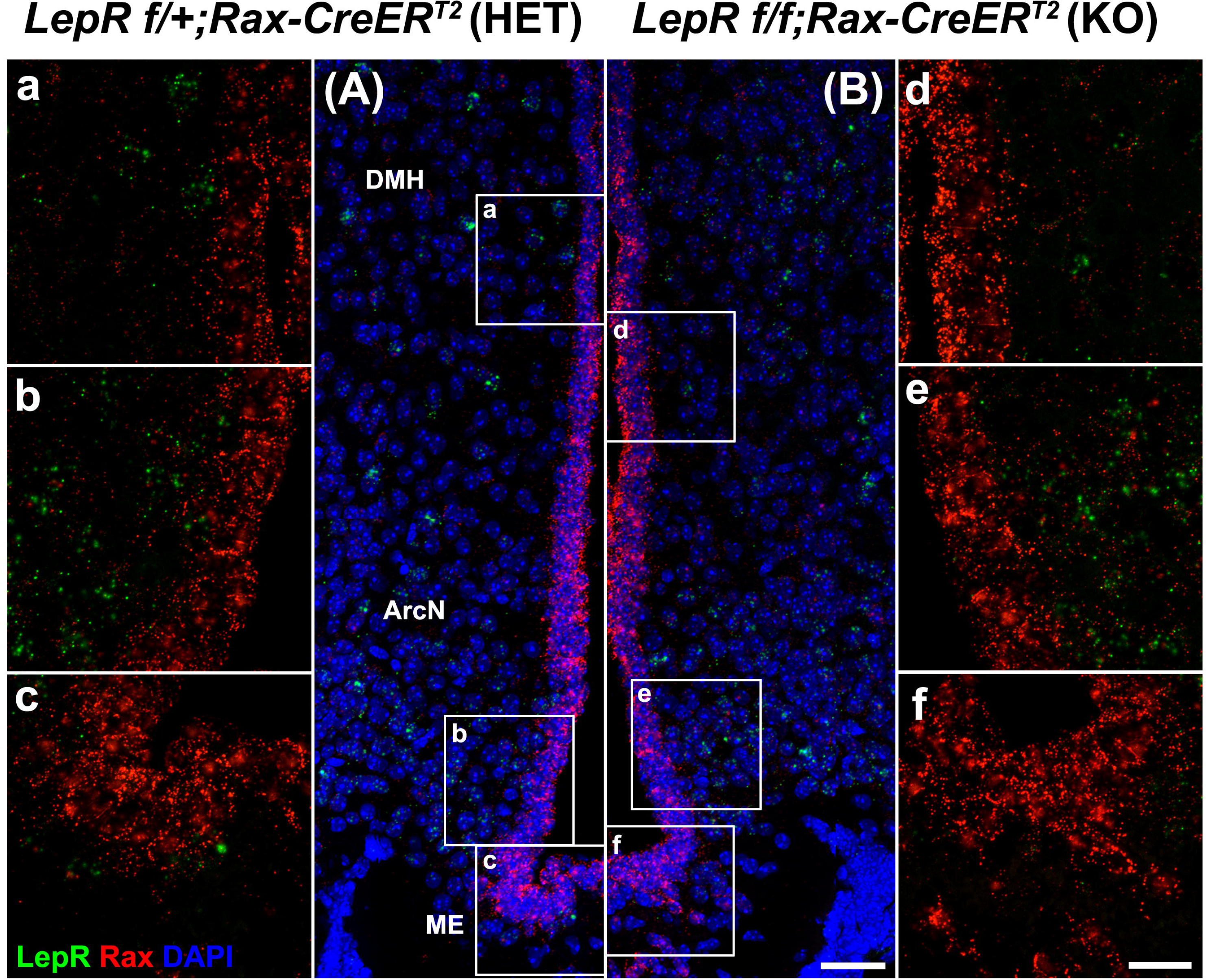
Tanycyte-specific genetic ablation of *LepR*. SmfISH analysis using *Rax*(red) and *LepR*(green) probes in *LepR^lox/+^;Rax-CreER^T2^* (A, HET) and *LepR^lox/lox^;Rax-CreER^T2^* (B, KO). (a-c) and (d-f) are the higher magnification images of the boxed area in (A) and (B), respectively. Scale bar: 50um (A, B), 20um (a-f)

We next examined the effect of tanycyte-specific *LepR* deletion on hypothalamic leptin response. Phosphorylation of signal transducer and activator of transcription 3 (STAT3) was used as readout of leptin signaling. Contrary to the previous reports (Balland et al., 2014), we did not observe pSTAT3 staining in tanycytic layer at 5 min after intraperitoneal (i.p.) leptin injection in both heterozygous *Rax-CreER^T2^;LepR^lox/+^;CAG-lsl-tdTom* and *LepR*-deficient *Rax-CreER^T2^;LepR^lox/lox^;R26-lsl-tdTom* mice (Figure 4A). By 45 min following treatment, pSTAT3-positive cells are observed throughout the hypothalamic parenchyma, including in the ventromedial nucleus (VMH), dorsomedial nucleus (DMH) and lateral hypothalamus (LH), but no difference in the number of immunopositive cells or the intensity of pSTAT3 staining was observed between heterozygous control and mutant mice (Figure 4C).

**Figure 4.**
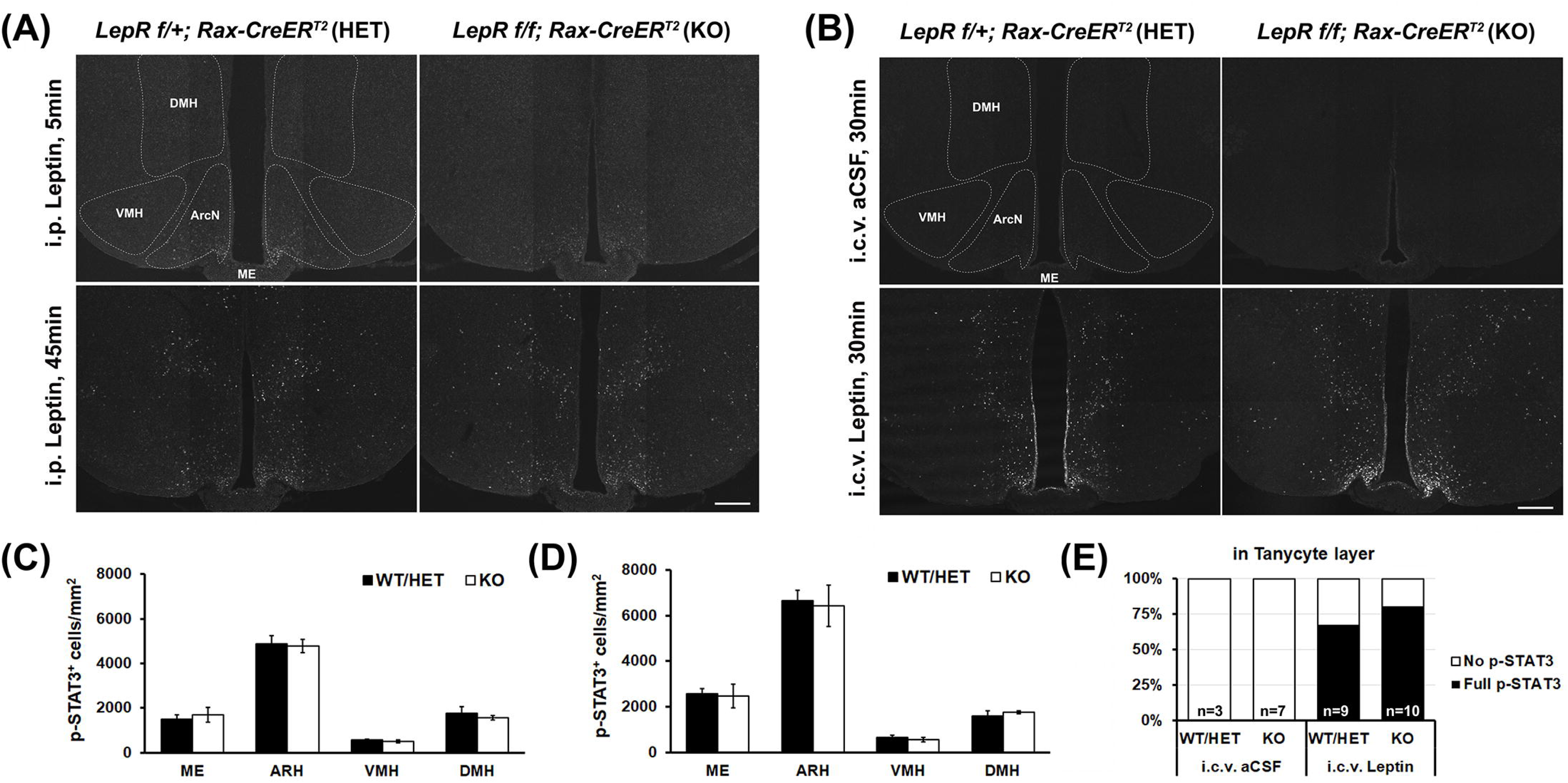
Leptin-induced STAT3 phosphorylation in the hypothalamus of control and *LepR* cKO mice. (A) Representative images of pSTAT3 immunohistochemistry 5min (upper) and 45min (lower) after i.p. leptin injection. (B) Quantification of the pSTAT3 positive cells in the images shown in (A), n=3-4. (C) Representative images of pSTAT3 immunohistochemistry 30min after i.c.v. aCSF (upper) and leptin (lower) injection. (D) Quantification of the pSTAT3 positive cells in the images shown in (C), n=3-4. (E) Percentage of cases showing the tanycytic pSTAT3 in each indicated condition. DMH, dorsomedial nucleus; VMH, ventromedial nucleus; ArcN, arcuate nucleus; ME, median eminence. Scale bar: 200um (A, C)

To investigate the effect of central leptin injection, either leptin or artificial cerebrospinal fluid (aCSF) was directly injected into the right lateral ventricle (LV) through an implanted cannula. 30 min following leptin, parenchymal neurons show the similar patterns of leptin-induced pSTAT3 staining, regardless of genotype (Figure 4B, D). In addition, considerable pSTAT3 signal is observed in the tanycytic layer, which is clearly distinct from ependymal cells. This paradoxical leptin-induced STAT3 phosphorylation in tanycytes that lack functional *LepR* implies the presence of an indirect, leptin-regulated mechanism that controls this process.

### Identification of leptin-regulated genes in ventrobasal hypothalamus using act-Seq

This observation of leptin-dependent, but LepR-independent, induction of pSTAT3 in tanycytes suggests two possibilities. First, that leptin can act on tanycytes directly via yet-to-be identified receptor(s) or via an indirect pathway that induces phosphorylation of STAT3. As no LepR-independent mechanisms of leptin signaling have yet been described, this is far more likely to reflect action of cytokine signaling triggered as a result of LepR signaling in other cell types, particularly endothelial cells and neurons. More generally, the genes regulated in response to leptin signaling in hypothalamus are not well characterized (Schwartz et al., 1996; Jovanovic et al., 2010). We sought to apply the newly developed technique of act-Seq to profile hypothalamic cells following i.c.v. delivery of leptin. This approach combines scRNA-Seq with actinomycin D treatment, which blocks *de novo* transcription following cell dissociation, and captures a snapshot of stimulus-induced changes in gene expression (Wu et al., 2017).

We conducted act-Seq analysis on the mediobasal hypothalamus one hour following a single i.c.v. infusion of either aCSF or leptin (Figure 5A). This interval was chosen to capture initial transcriptional changes induced following leptin treatment, soon after phosphorylation of STAT3 is detected. Using act-Seq, we can capture all main neuronal and non-neuronal populations in the brain (Figure 5A). Cells from aCSF-and leptin-treated samples aggregate together (Supplementary Figure 5A.), indicating that leptin infusion itself did not substantially change cell type-specific gene expression profiles. Clusters were identified based on enriched expression of previously reported marker genes (Supplementary Figure 1B, C), as well as previous scRNA-Seq studies of ventrobasal hypothalamus (Campbell et al., 2017; Chen et al., 2017). Individual clusters were extracted and expression were compared between treatment groups. Multiple genes are significantly differentially expressed in each cell cluster (Figure 5B-D, Supplementary Table 1).

**Figure 5.**
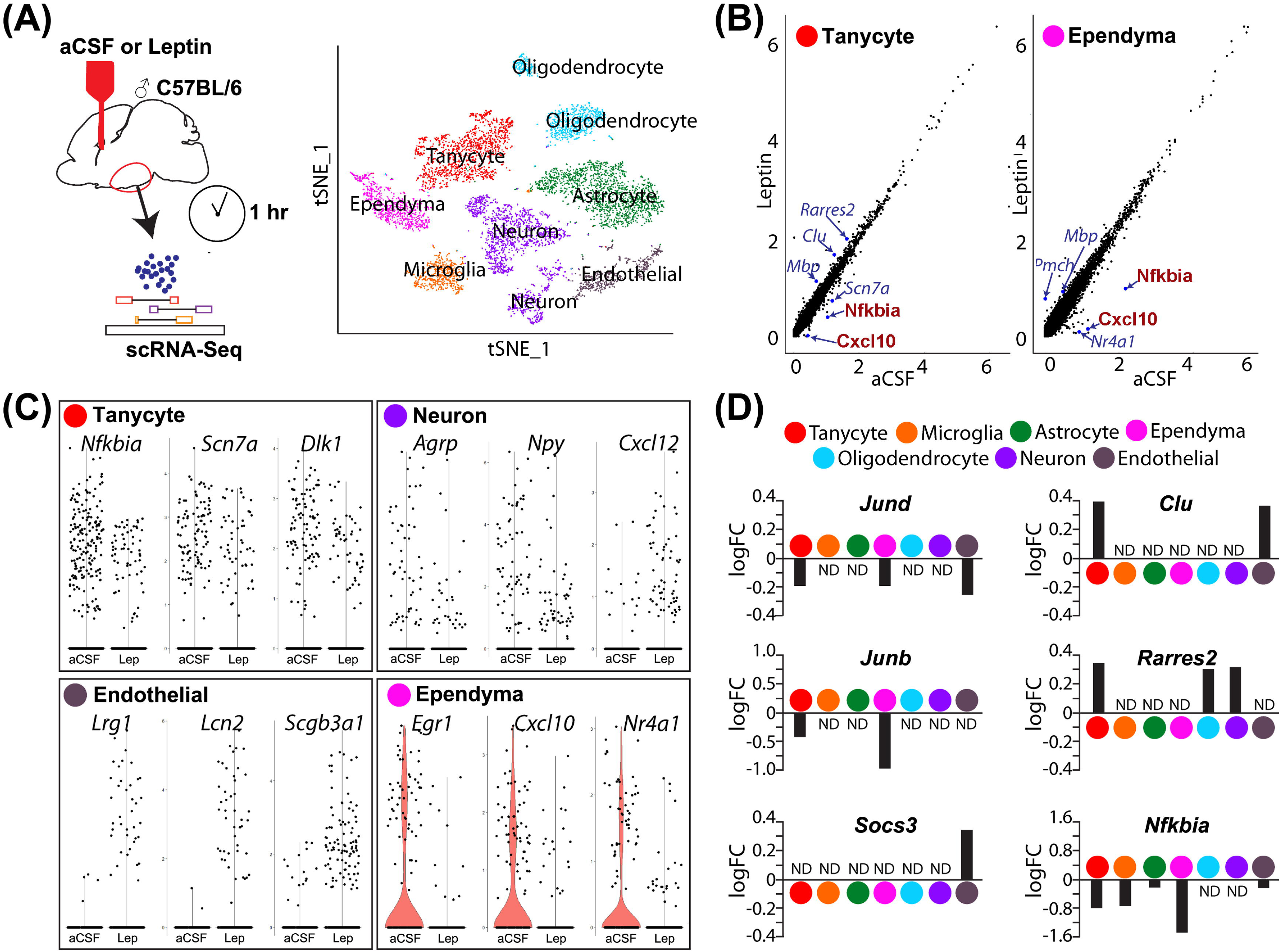
scRNA-Seq analysis identifies differentially expressed genes in mediobasal hypothalamus following i.c.v. leptin infusion. (A) Schematic diagram describing the pipeline of generating act-Seq data (left) and 2D tSNE plot with annotated clusters (right). (B) Correlation plot showing log-normalized gene-expressions between aCSF (x-axis)-and leptin (y-axis)-infused samples in tanycytes (left) and ependymocytes (rights). (C) Violin plots showing changes in gene expressions between aCSF-and leptin-infused samples in tanycytes (top left), neurons (top right), endothelial cells (bottom left), and ependymocytes (bottom right). (D) Bar graphs showing fold changes between two groups (positive fold change indicates an increased expression in leptin-infused group) in all 7 clusters.

Changes in gene expression are observed in all cell clusters following leptin infusion. The most noticeable change is the decreased expression of multiple *Nfkb* inhibitor family genes and chemokines in multiple cell clusters (Figure 5, Supplementary Table 1). For example, the inhibitor of *Nfkb* signaling *Nfkbia* and a chemokine of the C-X-C subfamily *Cxcl10* both show marked reductions in both tanycytes and ependymal cells (Figure 5B, D). The other differentially expressed genes substantially differ among cell types (Figure 5C, Supplementary Table 1). In tanycytes, leptin infusion significantly reduces expressions of the voltage-gated sodium channel *Scn7a* and the non-canonical Notch ligand *Dlk1* (Figure 5C). Leptin infusion decreases the level of *Agrp* and *Npy* expression in neurons, along with expression of the chemokine *Cxcl12* (Figure 5C). Expression levels of *Agrp* and *Npy* are known to be regulated by leptin in neurons (Korner et al., 2001; Morrison et al., 2005). *Socs3* antagonizes pSTAT3-dependent transcriptional activation following leptin (Howard et al., 2004; Mori et al., 2004; Morrison et al., 2005). *Socs3* levels are significantly upregulated in endothelial cells (Figure 5D), but not detectably induced in other cell types. Endothelial cells show a sharp increase in expression of the leucine-rich alpha 2 glycoprotein *Lrg1*, the Iron-trafficking protein *Lcn2*, and the secretoglobin *Scgb3a1* following central leptin infusion (Figure 5C).

Differential expression of immediate early genes is also detected in multiple clusters. The early growth response *Egr1 expression* was dramatically inhibited following leptin treatment in ependymal cells (Figure 5C). Expression of Jun family genes, *Jund* and *Junb*, was reduced in both tanycytes and ependymal cells (Figure. 5D). We also found that expression of the nuclear receptors *Nr4a1* and *Nr4a3* decreased substantially (Figure 5C, Supplementary Table 1). Tanycytes showed increased expression of *Clu* and *Rarres2* following leptin infusion, although this change was not detected in ependymal cells, where these genes are expressed at higher basal levels (Figure. 5D). With scRNA-Seq, we are also able to detect leptin-induced changes in gene expression oligodendrocytes and astrocytes (Figure 5C, D; Supplementary Table 1).

## Discussion

This study used multiple methods --qRT-PCR, smfISH, and both bulk and scRNA-Seq --to demonstrate that no detectable *LepR* expression is present in tanycytes. In addition, this absence, or very low level, of *LepR* expression in tanycytes is also seen in neonatal animals, and was not altered by fasting. Moreover, selective disruption of *LepR* in tanycytes does not affect levels of leptin-induced STAT3 phosphorylation in hypothalamic neurons, regardless of whether leptin was administered via an i.p. or i.c.v. route. Leptin-induced STAT3 phosphorylation was observed in tanycytes only following i.c.v. leptin delivery, suggesting that leptin may indirectly regulate tanycyte function. To better understand this process, we generated scRNA-Seq libraries from mice following i.c.v. leptin administration, and identify leptin-regulated genes, and possible mechanisms of crosstalk between tanycytes and other cell types, as discussed below.

### Previous reports of LepR expression in tanycytes

In this study, using multiple approaches, we fail to find evidence for expression of functional *LepR* in tanycytes. Our findings are in line with those reported recently by other groups. A study on *LepR* expression in non-neuronal brain cells using *ObRb-Cre* mice clearly demonstrated that the expression of fluorescent reporter was absent in the 3rd ventricular layer (DeFalco et al., 2001; Yuan et al., 2018). Consistent with this observation, several studies using the same mouse line failed to report *LepR* expression in this brain region (DeFalco et al., 2001; Scott et al., 2009; Djogo et al., 2016; Di Spiezio et al., 2018; Yuan et al., 2018). Likewise, similar results were found using a second *LepR-Cre* mouse reporter line (Leshan et al., 2006; Patterson et al., 2011).

The reasons for the discrepancy between our findings and previous studies reporting LepR expression in tanycytes remain unclear. Previous studies reporting an active role of LepR in leptin transport used qRT-PCR to measure *LepR* mRNA expression in tanycytes maintained in primary culture (Balland et al., 2014). Maintaining tanycytes *ex vivo* in this manner may have activated LepR transcription. A second study showed that *LepR* expression in the 3rd ventricle drops dramatically in the first postnatal week in rats with a sharp decrease in expression (Cottrell et al., 2009). We are unable to detect *LepR* expression in our study, even in neonatal mice (Figure 2B). It is possible that this may reflect a species-specific difference in gene expression between rats and mice. Although several other studies have reported leptin-triggered physiological responses in tanycytes and other cells of the hypothalamic ventricular layer (Hubschle et al., 2001; Frontini et al., 2008; Gil et al., 2013), we conclude that tanycytes and ependymal cells respond to the i.c.v.-infused leptin independent of *LepR*, and that these two cell types do not directly uptake leptin into brain and thereby *LepR* signaling in hypothalamic neurons.

### Indirect leptin-dependent regulation of gene expssion in tanycytes and ependymal cells

Any actions of leptin in tanycytes can be either direct, through *LepR*-independent signaling by leptin, or indirect, through signals released from other leptin-responsive cells. Our act-Seq data provides insights into the possible mechanisms by which leptin regulates tanycyte and ependymal cell function. *Lrp2* (low-density lipoprotein receptor-related protein-2, known as Megalin) has been known to bind and uptake leptin although it is also a potential receptor for multiple other secreted ligands, including clusterin (*Clu*) (Hama et al., 2004; Dietrich et al., 2008; Gil et al., 2013). ScRNA-Seq analysis shows that *Lrp2* is exclusively expressed by ependymal cells. Interestingly, *Clu* expression was significantly increased in both endothelial cells and tanycytes by central leptin infusion. Moreover, two recent studies show that *Il6r* and *Cntfr* expression are expressed in tanycytes, and activation of both receptors is sufficient to induce pSTAT3 in tanycytes (Severi et al., 2015; Anesten et al., 2017). However, it is not clear if either IL6 or CNTF release is modulated by leptin, or what the cellular origin of these ligands might be.

It is notable that emerging evidence supports that leptin resistance is unlikely to result from impaired transport of leptin across the BBB (Harrison et al., 2018; Kleinert et al., 2018). Leptin is more likely to be naturally transported into the brain through the *LepR* expressed in choroid plexus (CP), where it is very strongly expressed. Using high resolution imaging, these studies demonstrated that ependymal cells or α tanycytes do not directly take up leptin. However, the authors also observed an accumulation of fluorescent-labeled leptin in β2 tanycytes, consistent with a previous report (Balland et al., 2014). In this study, it is possible that leptin that is transported into the cerebral ventricles via the CP is trapped in the third ventricular floor through a *LepR*-independent mechanism. More importantly, the study did not show any correlation between this accumulation of leptin and the onset of leptin resistance.

### LepR-independent pSTAT3 signaling and anti-inflammatory effect of leptin in tanycytes

Previous studies reported that STAT3 phosphorylation induced by i.p leptin occurred more rapidly in tanycytes than in neurons (Balland et al., 2014). However, we were unable to reproduce this finding using the same methodology (Figure 4A). Instead, we observed strong activation of pSTAT3 simultaneously in all tanycyte subtypes only after i.c.v,. but not after i.p, leptin injection (Figure 4C). Leptin-induced pSTAT3 immunostaining has been described following i.c.v. injection in tanycytes in rats, where transient staining is observed beginning at 15 min following injection (Hubschle et al., 2001). However, the functional significance of this i.c.v leptin-induced pSTAT3 in tanycytes remains unknown, as is the identity of the ligand that induces it.

Tanycytes have long basal processes that extend into the hypothalamic parenchyma, which make this cell type both distinct from ependymal cells and well positioned to communicate with a large and diverse population of hypothalamic cell types. Ultrastructure studies have shown that tanycyte processes are found in close proximity with microvessels (Bleier, 1971; Peruzzo et al., 2004; Rodriguez et al., 2005; Miyata, 2015). Because of this unique structure and localization, tanycytes are likely to allow the vascular structure of this particular region of the brain to be more accessible and dynamic under certain physiological conditions. Classically, inflammatory cytokines have been implicated in the endothelial cell junctional disruption and vascular permeability, mostly in cancer research and recently in metabolic disease models (Graupera and Claret, 2018). Endothelial barrier integrity is also regulated by adhesion molecules (Vandenbroucke et al., 2008; Sukriti et al., 2014). Interestingly, our scRNA-Seq data showed a global reduction in expression of inflammatory factors and a number of junctional markers following i.c.v. leptin injection. We also observed a leptin-dependent down-regulation of activity-regulated immediate-early genes (IEG) --such as *Fos, Jun* and *Egr1* --in both ependymal cells and tanycytes, as well as endothelial cells and mediobasal hypothalamic neurons, which express *LepR* and are directly leptin-responsive (Elmquist et al., 1998; Mutze et al., 2006). It is unclear whether these reductions in inflammatory gene and IEG expression are functionally linked (Supplementary Table 1).

Tanycytes also exhibit a leptin-induced decrease in *Vegfa* expression, which is a growth factor inducing angiogenesis or permeabilization of blood vessel during inflammation (Croll et al., 2004; Kumar et al., 2009; Argaw et al., 2012; Shaik-Dasthagirisaheb et al., 2013). In line with our findings, *Vegfa* gene expression has been shown to increase in tanycytes during fasting, resulting in vascular remodeling in hypothalamic parenchyma (Langlet et al., 2013a). Tanycytic *Vegfa* promoted endothelial cell permeability in the ME, while increasing the expression of tanycytic tight junction proteins. However, since fasting has been linked with anti-inflammatory effect in the brain (Lavin et al., 2011), the observed reduction in *Vegfa* expression does not seem to correlate with inflammatory status. In our study, despite reduced *Vegfa* expression in tanycytes, leptin eventually led to loosening of the BBB by clear induction of the cytokine-like molecules, such as *Lcn2* and *Lrg1*, in endothelial cells (Lee et al., 2011; Wang et al., 2013; Jin et al., 2014). Future studies are thus warranted to find the signaling molecules that mediate communication between tanycytic processes and parenchymal microvessels. To assist with these studies, we provide a list of related genes showing significant differential expression in response to the i.c.v. injection (Supplementary Table 1).

---

**Supplementary Figure 1. Act-Seq analysis of leptin and saline-treated ventrobasal hypothalamus.**

(A) 2D tSNE plot showing distribution of two treatment groups. (B) Heatmap showing highly-expressed genes in all 7 cell populations in both treatment groups. (C) 2D tSNE plots showing annotated clusters in different colors (left) and enriched gene expressions between clusters (right).

**Supplementary Table 1. Genes differentially regulated in each mediobasal hypothalamic cell cluster following leptin treatment.**

All genes that showed differential expression between saline and leptin-treated samples are shown by cell type. The log_2_ fold-change is listed for leptin relative to saline-treated samples, along with the fraction of cells in the cluster showing expression of the gene in question in saline and leptin-treated samples, and both the raw and adjusted p-value for the comparison (calculated via Wilcoxon t-test).

## Acknowledgements

We thank W. Yap for comments on the manuscript. This work was supported by grant DK108230 to S.B. and a Maryland Stem Cell Postdoctoral Research Fellowship to S.Y. Bulk and scRNA-Seq data has been submitted to GEO.

