## Supplementary figures and images for "Tanycyte-independent control of hypothalamic leptin signaling"

### Supplemental Figure 1

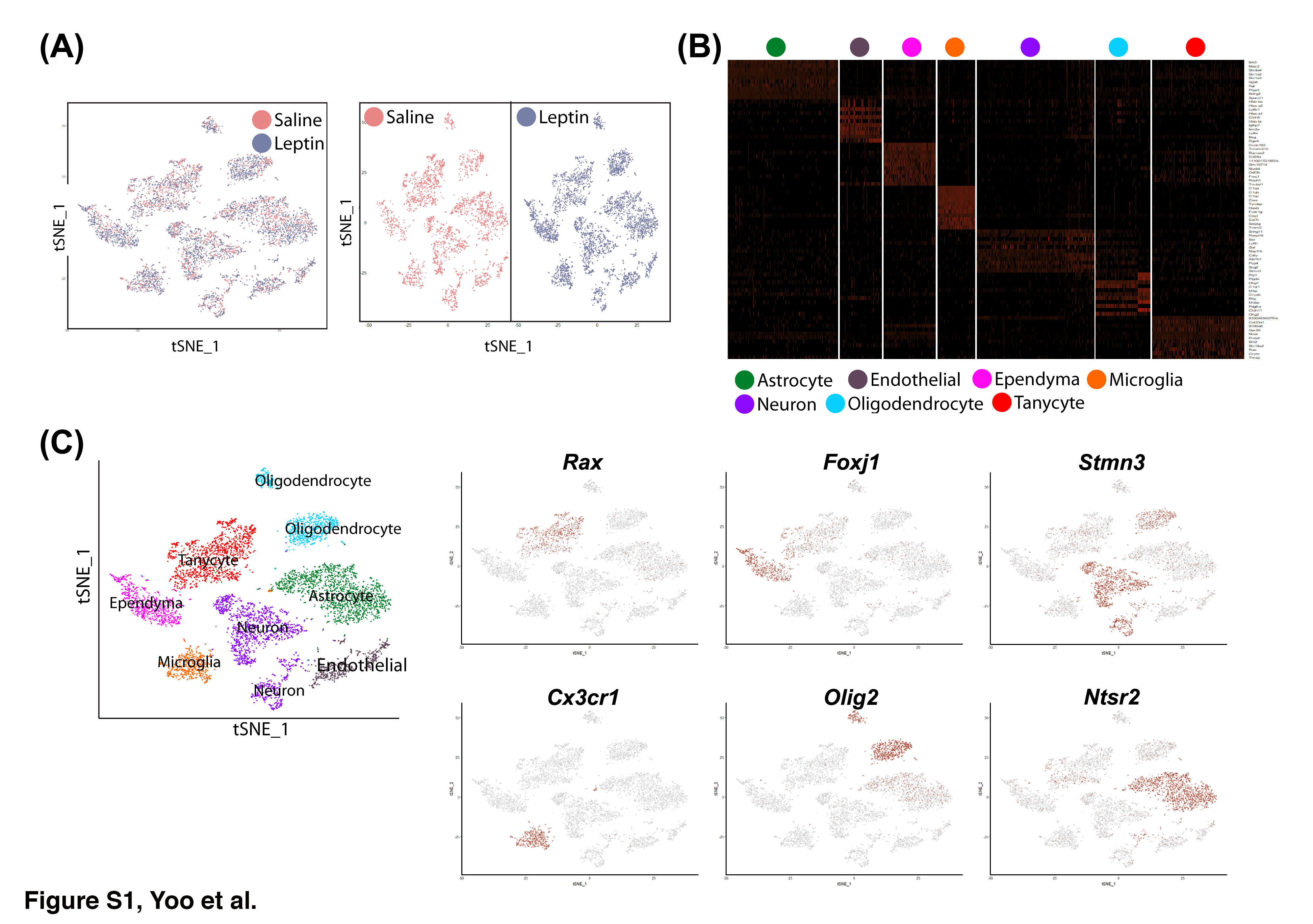
